# Efficient production of erythroid, megakaryoid and myeloid cells, using single cell-derived iPSC colony differentiation

**DOI:** 10.1101/246751

**Authors:** Marten Hansen, Eszter Varga, Cathelijn Aarts, Tatjana Wust, Taco Kuijpers, Marieke von Lindern, Emile van den Akker

## Abstract

Hematopoietic differentiation of human induced pluripotent stem cells (iPSCs) provide opportunities not only for fundamental research and disease modelling/drug testing but also for large-scale production of blood effector cells for future clinical application. Although there are multiple ways to differentiate human iPSCs towards hematopoietic lineages, there is a need to develop reproducible and robust protocols. Here we introduce an efficient way to produce three major blood cell types using a standardized differentiation protocol that starts with a single hematopoietic initiation step. This system is feeder-free, avoids EB-formation, starts with a hematopoietic initiation step based on a novel single cell-derived iPSC colony differentiation and produces multi-potential progenitors within 8-10 days. Followed by lineage-specific growth factor supplementation these cells can be matured into well characterized erythroid, megakaryoid and myeloid cells with high-purity, without transcription factor overexpression or any kind of pre-purification step. This standardized differentiation system provides a simple platform to produce specific blood cells in a reproducible manner for hematopoietic development studies, disease modelling, drug testing and the potential for future therapeutic applications.

**Highlights:**

- Efficient hematopoietic differentiation from single cell-derived iPSC colonies
- Reproducible feeder-free, monolayer differentiation system independent of iPSC line
- Production of erythroid, megakaryoid and myeloid cells with high-purity
- Platform for hematopoietic developmental research and future clinical application

## Introduction

The discovery that human somatic cells can be reprogrammed into a pluripotent stem cell state, has created a novel approach for developmental studies, drug screening/discovery, disease modelling, and regenerative medicine (Takahashi et al, 2007; Takahashi & Yamanaka, 2006; Wu & Hochedlinger, 2011). Recent advances in hematopoietic differentiation of human induced pluripotent stem cells (iPSCs) opened opportunities not only for fundamental research and disease modelling/drug testing but also for large-scale manufacture of blood products with the expectancy of clinical application (Giarratana et al, 2011; Ross & Akimov, 2014). However, this potential is influenced by the I) variations at (epi)genetic level between iPSC lines or between clones, II) incomplete understanding of differentiation cues, III) lack of standard (GMP-grade) protocols, IV) low cellular yield and V) low purity, often requires pre-purification. These also make direct comparisons between the final results obtained in the different laboratories difficult. Thus, there is a need of improvement and standardization of iPSCs differentiation to hematopoietic cells in order to obtain a reproducible method which yields cells enough for downstream applications. In contrast to mouse embryonic stem cells (ESCs)/iPSCs, human iPSC are sensitive for single cell-passaging due to detachment-induced apoptosis (Amit & Itskovitz-Eldor, 2002; Chen et al, 2010; Ohgushi et al, 2010). Therefore, clump-passaging is still the preferred method for human ESC or iPSC maintenance (Hartung et al, 2010; Reubinoff et al, 2000). Besides its labour intensiveness, clump-passaging technically leads to heterogeneity in clump size, causing potential reproducibility issues. Dissociation-induced apoptosis of iPSCs is mediated by the Rho-associated protein kinases (ROCK) pathway (Chen et al, 2010; Ohgushi et al, 2010; Watanabe et al, 2007). The use of ROCK inhibitors during single cell-dissociation significantly increases cell survival, colony formation and decreases cell death without affecting pluripotency, thereby enabling single cell-passaging (Kajabadi et al, 2015; Kurosawa, 2012; Watanabe et al, 2007; Zhang et al, 2016).

Differentiation of iPSCs towards the hematopoietic lineage can be achieved by 2D- or 3D methods. Forced or spontaneous clustering of iPSCs into a spherical 3D structures (embryoid body (EB)) recapitulates in some aspects the complex assembly during embryogenesis (Amit & Itskovitz-Eldor, 2002; Doetschman et al, 1985; Ng et al, 2005), as evidenced by the formation of the three germ layers (Amit & Itskovitz-Eldor, 2002; Itskovitz-Eldor et al, 2000). Differentiation can be directed towards cell types of interest by controlling the EB size, or by using extracellular matrix and/or growth factors. With the development of forced aggregation, EB size can be better controlled but spontaneous agglomeration of multiple EBs is still an unregulated factor (Ng et al, 2008). Release of progenitors from the EBs involves either attaching EBs to a matrix coated surface or applying physical shear stress sometimes combined with the use of proteases (Fujita et al, 2016; Ko et al, 2014; Olivier et al, 2016). These are harsh treatments, affecting viability and the ability to further culture these cells. As an alternative to the 3D approach, iPSCs can be differentiated in 2D monolayers on a feeder cell layer or on feeder-free matrix coated dishes (Choi et al, 2009a; Choi et al, 2009b; Feng et al, 2014; Niwa et al, 2011; Salvagiotto et al, 2011; Smith et al, 2013; Weisel et al, 2006). The 2D method enables the collection of hematopoietic cells without the use of proteases or physical shear stress, increasing the yield of viable cells (Feng et al, 2014; Niwa et al, 2011; Salvagiotto et al, 2011). Currently, 2D protocols have been initiated using iPSC clumps of variable size, hindering reproducibility and robustness. We hypothesized that tight control of iPSC differentiation starting conditions will increase the reproducibility of hematopoietic specification and differentiation.

In the present study a novel, feeder-free, single iPSC-derived, monolayer differentiation system was designed to culture 3 major myeloid blood cell types starting from a single hematopoietic initiation step. The derived multi-potential hematopoietic progenitors could further mature into erythroid, megakaryoid and myeloid cells with high purity. This protocol provides a reproducible platform not only for hematopoietic developmental research but for disease modelling/drug testing and it has the potential for large-scale production of specific blood cell types for future clinical application.

## Material and methods

### Chemicals

The chemicals were purchased from Sigma-Aldrich (Munich, Germany), the culture reagents from Thermo Fisher Scientific (Waltham, Massachusetts, USA) and all growth factors from Peprotech (Pittsburgh, USA) unless otherwise specified.

### Statement

Informed consent was given in accordance with the Declaration of Helsinki and Dutch national and Sanquin internal ethic boards.

### iPSC lines

Four iPSC lines were used in this study. MML-6838-Cl2: megakaryoblast origin, hOKSM lenti-virus vector; MEL1171.CL7: erythroblast origin, hOKSM lenti-virus vector; SANi003-B: erythroblast origin, hOKSML episomal plasmid (transgene-free), SANi003-A: erythroblast origin, hOKSML episomal plasmid (transgene-free) (Hansen et al, 2017; Varga et al, 2017). All used lines are registered at the human pluripotent stem cell registry (https://hpscreg.eu).

### iPSC single cell-maintenance

All cells were maintained at 37°C in humidified atmosphere containing 5% CO_2_. The iPSCs were cultured feeder-free on matrigel (BD Biosciences) in essential 8 medium (E8). Passaging occurred every 5-7 days using TrypLE Select according to manufacturer’s instruction, with a seeding density of 1.0-2*10^4^ cells/6 cm dish in E8 + Revitacell supplement. If necessary, differentiated parts were removed by scraping. Cells were counted on a Casy cell counter and analyser (Roche).

### iPSC colony differentiation towards hematopoietic lineages

In order to reach the required colony number (5-15 colonies/6 cm dish) the cells were harvested as described above. The iPSCs were single cell-seeded at lower density, 250-400 cells/6 cm dish on matrigel (BD Biosciences) coated dishes. Single cell-derived colonies were grown for 8-9 days prior to differentiation. If necessary, surplus colonies were removed by scraping. At day 0 of differentiation, cells were washed 1x with DPBS then hematopoietic differentiation was initiated by adding Stemline II supplemented with 1% Pen/strep, 50 ng/ml bFGF, 40 ng/ml VEGF, 20 ng/ml BMP4 and 100x insulin-transferrin-selenium (ITS). At day 6, colonies were washed 1x with DPBS then culture media was added containing lineage specific growth factor cocktails (described below) and re-freshed every 2-3 days.

### Specification towards erythrocytes

After initiation of differentiation for 6 days medium was changed to Cell-Quin, 100x ITS, 5 U/ml EPO, 50 ng/ml hSCF, 10 ng/ml TPO, 50 ng/ml IL-6, 50 ng/ml IL-3, 1% Pen/strep. Between day 9-15 of iPSC differentiation (day 9, 12, 15) suspension cell were harvested and analysed. Day 12 suspension cells were collected and further expanded for 9 days (day 12+9) in erythroblast (EBL)-specific media (Cell-Quin supplemented with: 1% Pen/strep, 2 U/ml EPO, 100 ng/ml hSCF, 10^−6^ M Dexamethasone)(Heideveld et al, 2015; van den Akker et al, 2010). Expanded iPSC-EBLs were further maturated by adding terminal differentiation mix Cell-Quin suppl. with: 1% Pen/strep, 10 U/ml EPO, 5 U/ml heparin, 5% pooled human plasma. 1*10^7^ cells were collected and analysed for Hb isoform expression by high-performance cation-exchange liquid chromatography (HPLC) on Waters Alliance 2690 equipment and the column was purchased from PolyLC (van Zwieten et al, 2014).

### Specification towards megakaryocytes

After 6 days of differentiation, medium was changed to Cell-Quin with 10 ng/ml VEGF, 20 ng/ml BMP4, 1 ng/ml IL-3, 10 ng/ml IL-6, 50 ng/ml TPO, 50 ng/ml hSCF, 100x ITS, 1% Pen/strep and 10 ng/ml IL-1β. Suspension cells were harvested between day 11-15 of iPSC differentiation and terminally differentiated in Cell-Quin with 1% Pen/strep, 10 ng/ml IL-1β and 50 ng/ml TPO for 6 days (day 11+6). For megakaryocyte colony assays (CFU-Mk) day 11 of iPSC differentiation, suspension cells were expanded for 6 days in Cell-Quin supplemented with 10 ng/ml IL-6, 50 ng/ml hSCF, 50 ng/ml TPO and 10 ng/ml IL-1β. The expanded cells were seeded into the Mk-CFU assay chamber slide (Stem cell technologies) at 5000 cells/well. Mk-CFU were fixed and stained according to manufactures instructions after 10 days of culture. For CD34^+^ mobilized peripheral blood (MPB) differentiation towards the megakaryoid linage, cells were differentiated in Cell-Quin with 50 ng/ml hSCF, 1 ng/ml IL-3, 20 ng/ml IL-6, 50 ng/ml TPO for 4 days followed by 10 days differentiation in Cell-Quin supplemented with 50 ng/ml TPO and 10 ng/ml IL-1β. Isolation of CD34^+^ MPB was performed as described before (Heideveld et al, 2015).

### Specification towards myeloid cells

From day 6 of differentiation, cells were cultured in Stemline II with 1% Pen/strep, 10 ng/ml VEGF, 20 ng/ml BMP4, 1 ng/ml IL-3, 10 ng/ml IL-6, 50 ng/ml TPO, 50 ng/ml hSCF, 100× ITS and 50 ng/ml FLT-3. Suspension cells were collected on day 12 of iPSC differentiation and further expanded for 3 days (day 12+3) in Stemline II supplemented with 1% Pen/strep, 10 ng/ml IL-3, 10 ng/ml GM-GSF, 30 ng/ml G-CSF, 50 ng/ml FLT-3, 50 ng/ml hSCF and after in Stemline II with 1% Pen/strep, 30 ng/ml G-CSF until day 12+7.

### Flow cytometry (FACS)

Cells were washed with DPBS, re-suspended in FACS buffer (1x DPBS, 0.1% BSA, 0.5 mM EDTA) and incubated with antibodies for 30 minutes at RT (Table 1, Table S1). Then cells were washed 2x with FACS buffer and the surface marker expression level was assessed by the LSR-II (BD Bioscience) or by the Canto-II (BD Bioscience). Data was analysed using Flowjo software.

**Table 1.** Used antibodies in this study.

| Antibodies | Manufacture | Dilution |
| --- | --- | --- |
| Anti CD34 – PB | Biolegend | 1:400 |
| Anti CD34 – APC | Biolegend | 1:500 |
| Anti CD41a – PE-Cy7 | Biolegend | 1:500 |
| Anti CD42b - FITC | Sanquin | 1:15 |
| Anti CD43 – PE-Cy7 | Biolegend | 1:200 |
| Anti CD36 – FITC | Pelicluster | 1:100 |
| Anti CD71 – APC | MACS | 1:200 |
| Anti CD71 – PB | MACS | 1:200 |
| Anti CD235a - PE | Acris | 1:2500 |
| Anti CD15 – FITC | BD | 1:150 |
| Anti CD11b – APC | BD | 1:1000 |
| Anti SIGLEC8 – PE | Biolegend | 1:100 |
| Anti CD123 – PE | BD | 1:10 |
| Anti CD18 – FITC | BD | 1:50 |
| Anti EMR3 - FITC | AbD Serotec | 1:100 |

### Image stream

Day 11+7 iPSC-megakaryoblasts (iPSC-MK) cells were stained for megakaryoid marker combinations CD34, CD41a and CD42a analysed with ImageStream^×^ Mark II (Merck). Size difference was calculated between population of megakaryoblast (CD34^+^/CD41a^+^) and megakaryocytes (CD41a^+^/CD42b^+^) with ImageJ.

### Statistical tests

Statistics were performed with GraphPad Prism, all quantifications are shown with SD and mean.

### Cytospins and staining

1*10^5^ cells were cytospun (Shandon CytoSpin II Cytocentrifuge) onto 76*26 mm glass microscope slides. The slides were air-dried and stained. Giemsa/benzidine staining was performed as previously described. For Giemsa May-Grünwald staining cytospins slides were incubated for 5 min in May-Grünwald followed by 15 min in phosphate buffer and subsequent staining Giemsa solution for 30 min. For qualitative peroxidase-activity cytospins were directly fixed after preparation in 10% formaldehyde vapours for 15 min at RT. After fixation, 2 drops of benzidine solution (benzidine in PBS + 0.1% H_2_O_2_) were added on the cell-spot and fixed cytospins were incubated for 30 min at 37°C. All slides were rinsed in deionized water, air-dried and analysed with Zeiss Scope.A1 microscope.

### Neutrophils isolated from blood (PMN)

Granulocytes were isolated by density gradient centrifugation with isotonic Percoll (Pharmacia). The pellet fraction, containing erythrocytes and granulocytes underwent erythrocyte lysis with ice-cold isotonic NH_4_Cl solution (155 mM NH_4_Cl, 10 mM KHCO_3_, 0.1 mM EDTA, pH 7.4) and were washed then re-suspended in Hepes-buffered saline solution (HBSS containing 132 mM NaCl, 6 mM KCl, 1 mM CaCl_2_ 1 mM MgSO_4_, 1.2 mM potassium phosphate, 20 mM Hepes, 5.5 mM, glucose and 0.5% (w/v) human serum albumin, pH 7.4)(Roos & de Boer, 1986). Functional assays of PMN and iPSC-derived myeloid cell were performed in Hepes-buffered saline solution.

### NADPH-oxidase and ROS production

NADPH-oxidase activity was assessed by the release of hydrogen peroxide, determined by an Amplex Red kit (Molecular Probes). Neutrophils and iPSC-derived granulocytes were left unstimulated or stimulated with phorbol-12-myristate-13-acetate (PMA, 100 ng/ml), in the presence of Amplex Red (0.5 μM) and horseradish peroxidase (1 U/ml). Fluorescence was measured at 30 second intervals for 20 min with the HTS7000plus plate reader (Tecan). Maximal slope of H_2_O_2_ release was assessed over a 2 min interval (Drewniak et al, 2010).

## Results

### Single cell-seeding allows the expansion of iPSC colonies whilst maintaining pluripotency

A robust and reproducible iPSC differentiation protocol should start with a defined range of iPSC colonies/surface that are synchronized in size. This was achieved by single cell-seeding with Revitacell that contains a more specific ROCK inhibitor than the commonly used thiazovinin and Y-27632. The morphology and pluripotency marker expression (SOX2, TRA-1-60, TRA-1-81, OCT4, SSEA4) of the iPSCs were unaffected during multiple single cell-passages using RevitaCell (Fig.1A, B and S1). In addition, iPSC lines retained their spontaneous differentiation ability towards all the 3 germ layers and showed a normal karyotype (Hansen et al, 2017; Varga et al, 2017). For differentiation, cells were seeded at 250-400 cells/6 cm dish, which gave rise to 5-15 colonies in a reproducible manner. iPSC colonies reached a size of approximately 300-600 μm within 8-9 days whilst maintaining proper iPSC morphology (Fig.1C and 2B). We observed size differences in colonies between iPSC lines (300-600 μm), within lines however there were minimal variations in size. These results indicate that uniform size and number of colonies can be generated from single cell-seeded iPSC, regardless of the iPSC line that was used.

**Figure 1.**
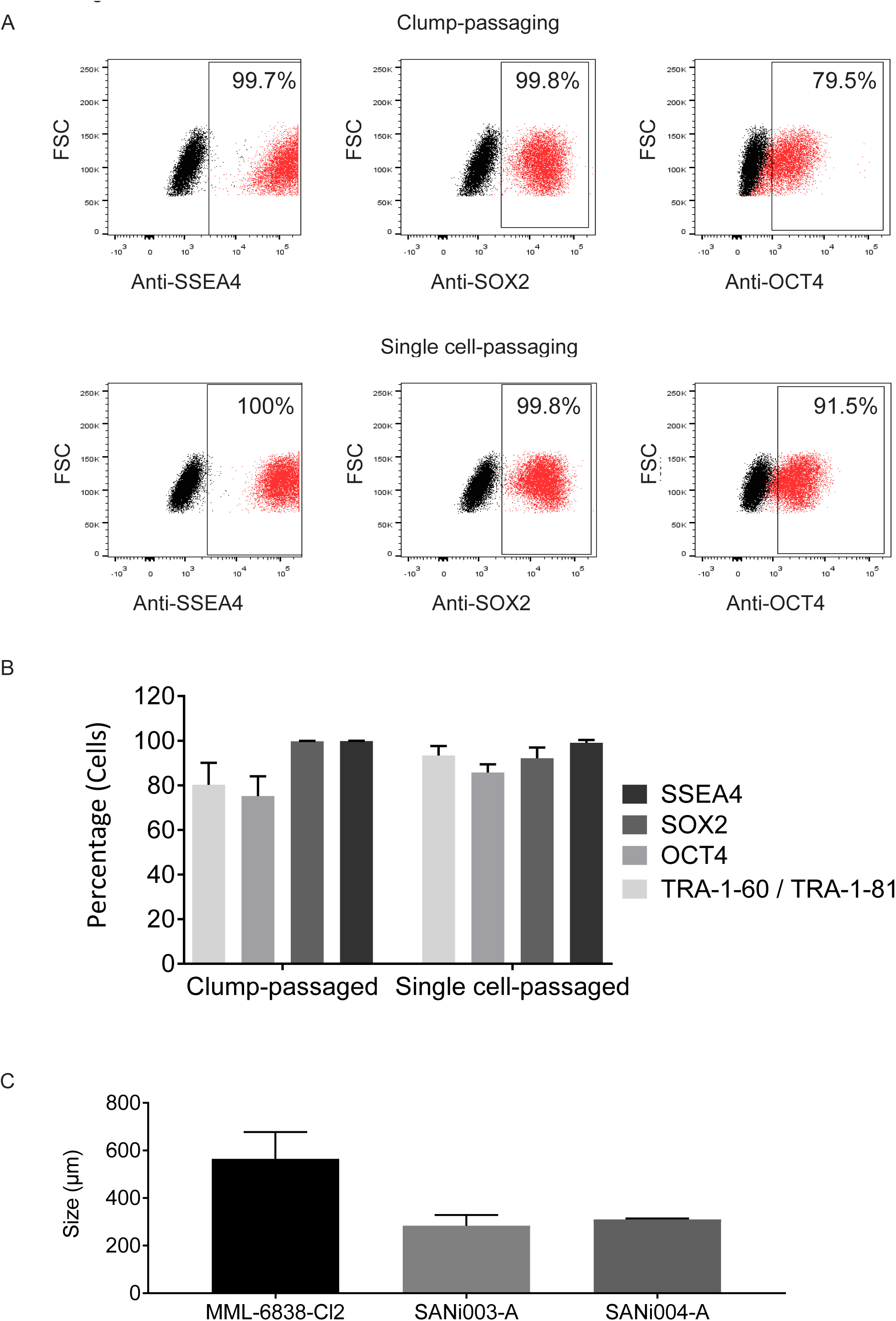
Pluripotency potential and colony size of single cell-seeded iPSCs. See also Figure S1. **A)** Clump-passaged and single cell-passaged iPSC were stained for pluripotency markers SSEA4, OCT4, SOX2 and analysed by flow cytometry (red: staining, black: isotype). **B)** The frequency (%) of pluripotency marker expression (SSEA4, OCT4, SOX2, TRA-1-60/TRA-1-81) was quantified (mean and SD for n=3) **C)** Quantification of colony size (μm) of the used iPSC lines 8-9 days after seeding, 6 measurements/clone, showing mean and SD, (n=3/clone).

**Figure 2.**
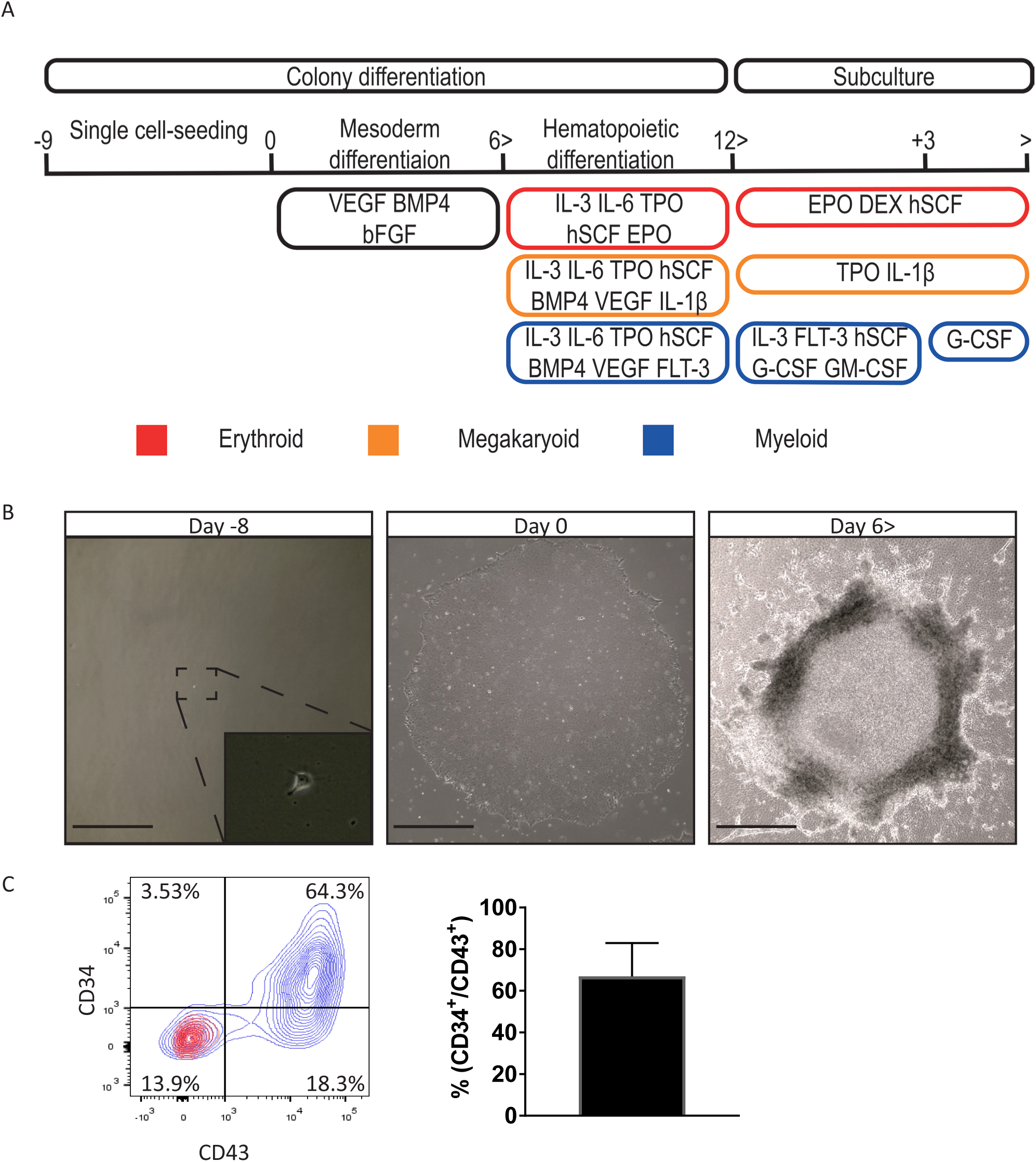
Direct differentiation of iPSC colonies to hematopoietic lineages. **A)** Schematic representation of single cell-seeded iPSC colony differentiation towards hematopoietic lineages, then further maturation of suspension cell to erythroid- (red), megakaryocyte- (yellow), myeloid cells (blue). **B)** Morphological changes of single cell-derived iPSC differentiation demonstrated by bright field microscopy (scale bar 200 μm). **C)** The appearance of CD34^+^/CD43^+^ population representing hematopoietic potential by flow cytometry (left, blue: staining, red: isotype) and quantification (right, n=5), showing mean and SD, “n” is biological replicates.

### Directed differentiation of iPSC colonies to hematopoietic lineages

Hematopoietic specification of iPSC follows embryonic development in some degree and involves induction of mesoderm, followed by a transition wave of endothelial to hematopoietic cell differentiation and the emergence of hematopoietic progenitors (Lim et al, 2013; Slukvin, 2013). We assessed differentiation of single cell-derived iPSC colonies in serum free media towards hematopoietic lineages by harvesting non adherent cells from the colony culture following the standardized hematopoietic induction scheme depicted in figure 2A. iPSC differentiation resulted in morphological changes within the colony and the appearance of CD43^+^/CD34^+^ cells that are associated with hematopoietic potential (Fig.2B,C) (Vodyanik et al, 2006). When the CD43^+^/CD34^+^ population emerged, the growth factor cocktail was changed by including basic hematopoietic growth factors (IL-6, IL-3, hSCF, TPO) and lineage specific growth factors of the cell type of interest: EPO (erythroid), IL-1β (megakaryoid) or FLT-3 (myeloid) (Fig.2A). Subsequently, hematopoietic suspension cells can be harvested between day 10-15. The quality, quantity and efficiency of differentiation potential was evaluated towards the erythroid, megakaryoid and myeloid lineages.

### iPSC differentiation to erythroid cells

The harvested non-adherent cells were directed to the erythroid lineage in serum-free medium as indicated in figure 2A. The purity of the erythroid population, and its distribution over different maturation stages was assessed by expression of CD36, CD71 (Transferrin receptor protein 1) and CD235a (Glycophorin A), using three different iPSC lines (MML-6838-Cl2; SANi003-A; SANi003-B)(Hansen et al, 2017; Varga et al, 2017). CD36^+^ in combination with CD71^+^/CD235a^+^ defines earlier erythroid population while CD36^-^/CD71^+^/CD235a^+^ represent a more differentiated erythroid population. Downregulation of CD36, followed by decreasing CD71 expression is a hallmark of EBL differentiation terminating in reticulocyte formation which is associated with CD71^low/-^ and CD235a^+^ expression (Li et al, 2014; Socolovsky et al, 2001). From day 9 of iPSC differentiation a non-adherent cell population was observed which was harvested until day 15. These suspension cells irrespective of collection day (day 9, 12, 15; from each iPSC clone) were highly enriched for erythroid cells: CD71^+^/CD235a^+^ (83,8%) and 64.4% CD36^+^ defining the early erythroid population. In addition, a small population of CD34^+^ (10.7%) was observed (Fig.3A and S2). These cells were negative for CD18 (myeloid lineage marker) confirming specification towards the erythroid lineage (Fig.3A). The harvested cells (day 12) could be further expanded for 9 days (day 12+9) in EBL expansion media containing EPO, hSCF and glucocorticoids (Fig.3B) (Leberbauer et al, 2005; van den Akker et al, 2010). On average, a single iPSC gave rise to ~8*10^3^ cells within 12 days and following 9 days expansion resulted in ~2*10^5^ erythroid cells/iPSC. During the erythroid expansion phase the CD36 expression decreased to 34%, while the cells reached 98.5% CD71^+^/CD235a^+^ purity (Fig.3B). The iPSC-derived CD71^+^/CD235a^+^, expanded erythroid cells (day 12+9) morphologically resembled basophilic and pro-erythroblast stages (Fig.3B). The expanded, pure iPSC-EBLs (day 12+9) were terminally differentiated to evaluate their maturation potential (Leberbauer et al, 2005; van den Akker et al, 2010). Terminal differentiation for 7 days gave rise to an erythroid population which was CD36^low^ (0.7%) but highly double positive for CD71/CD235a (90.6%). In addition, CD71^-^/CD235a^+^ expression was observed (7%) associated with reticulocyte formation (Fig.3C). The differentiated erythroid cells morphologically and phenotypically resembled a uniform late orthochromatic normoblast stage (Fig.3D). HPLC revealed that these matured erythroid cells expressed mainly α_2_-γ_2_ (fetal) hemoglobin (50.1%) and small amount of α_2_-β_2_ (adult) hemoglobin (2.1%), indicative of definitive fetal erythropoiesis (Fig.3D). In addition, we also found 15% of embryonic hemoglobin, indicating a low amount of primitive erythropoiesis (Fig.3D). The results overall showed that single cell-derived iPSC differentiation directed to erythroid cells successfully gave rise to an expandable, pure EBL population that can be further matured to orthochromatic normoblasts. Hemoglobin (Hb) subunit analysis corresponds to primarily definitive fetal erythropoiesis. Three different iPSC lines were used to validate the robustness of this differentiation method. The protocol was further tested in its potential to differentiate towards the megakaryocytic and myeloid lineages.

**Figure 3.**
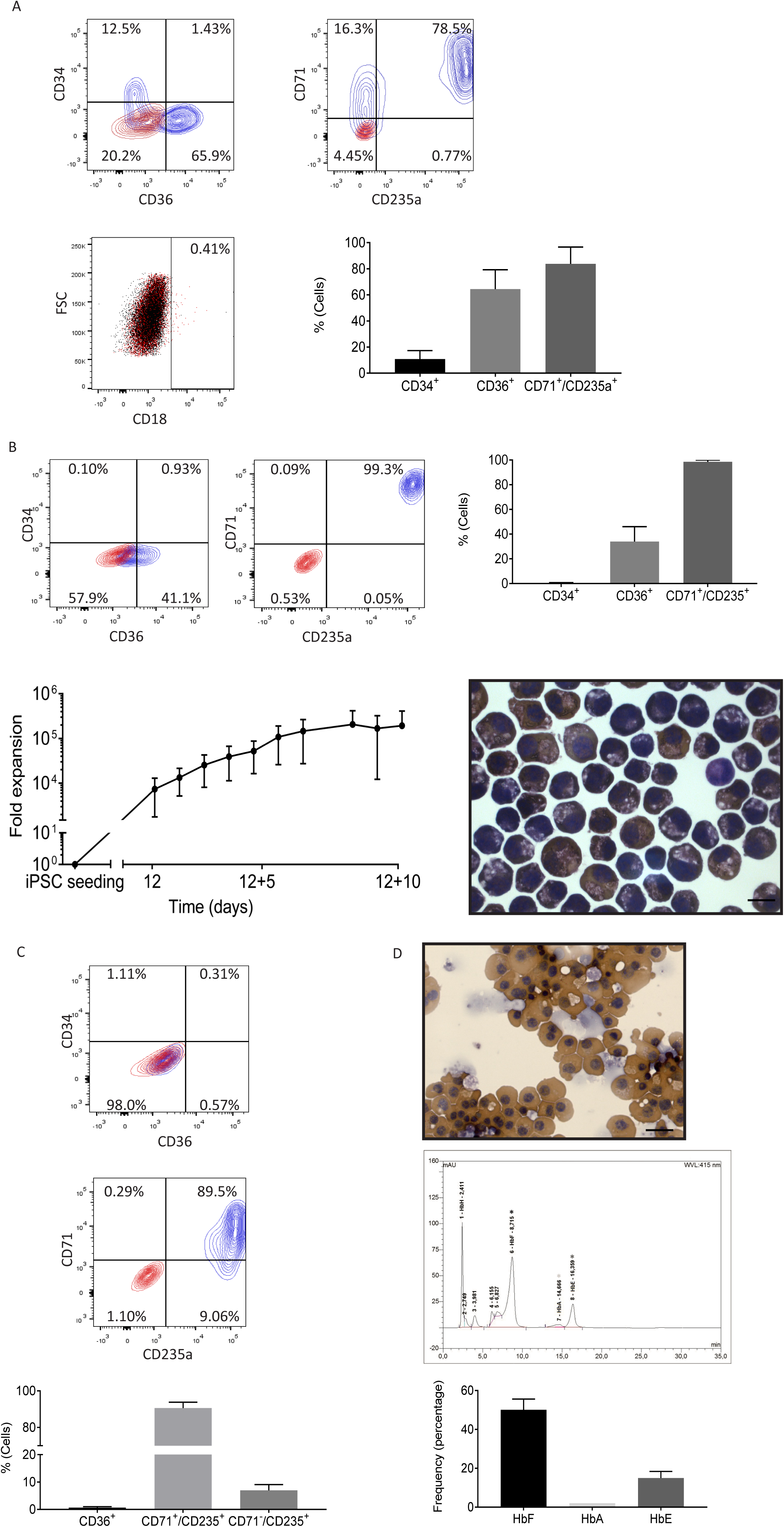
iPSC differentiation to erythroid cells. See also Figure S2. **A)** Erythroid-associated marker expression profile of suspension cells. Top: representative FACS plots (day 12) (blue: staining, red: isotype). Bottom-left: representative FACS plots showing the lack of CD18 expression (red: staining, black: isotype). Bottom-right: FACS quantification graph represents average expression of day 9, day 12, day 15 suspension cells (3 time points, 3 lines: n=9). **B)** Expression profile of expanded suspension cells for 9 days (day 12+9) demonstrated by FACS (top). Top-left: Representative FACS plots (blue: staining, red: isotype). Top-right: FACS quantification graph represents average expression (n=3). Bottom-left: growth curve of expanded suspension cells (n=3). Bottom-right: representative cytospin of expanded erythroid cells (scale bar 20 μm) **C)** Expression profile of erythroid cells after 7 days terminal differentiation demonstrated by FACS. Top and middle: Representative FACS plots (blue: staining, red: isotype). Bottom: FACS quantification graph represents average expression (n=3). **D)** Representative cytospin of erythroid cells after 7 days terminal differentiation (scale bar 20 μm) (Top). Representative HPLC measurement (Middle). Bottom: averaged % of adult-, fetal-, embryonic Hb (n=2). HPLC measurement shows mean, all other graphs show mean and SD, “n” is biological replicates.

### iPSC differentiation to megakaryocyte cells

Following hematopoietic commitment in serum-free media with the presence of TPO (figure 2A), a mixture of non-adherent megakaryoid cells, more immature progenitors and other cells were observed on day 11 of differentiation. The vast majority of this population was composed of megakaryoid cells at different stages (74.6%), including 25.2% megakaryoblasts (CD34^+^/CD41a^+^(ITGA2B)/CD42^-^(GP1B)), 14.6% late megakaryoblasts (CD34^-^/CD41a^+^/CD42^-^), 34.8% megakaryocytes (CD41a^+^/CD42^+^/CD34-). Non-megakaryoid cells included, CD34^+^/CD41a^-^/CD42^-^ cells (11.94%), likely a mix of hematopoietic progenitors and endothelial cells and 13.5% that were negative for all tested markers (CD34^-^/CD41a^-^/CD42^-^) (Fig.4A). This mix of mainly megakaryoid cells was further cultured for 6 days in the presence of TPO and IL1β, and subjected to megakaryoid colony assays (CFU-Mk). On average iPSC-MK gave rise to 102 (68-141) CFU-Mk/5000 cells, which is considerably higher than reference values for bone marrow mononucleated cells that produce 5 CFU-Mk/5000 cells on average (Fig.4B) (Hogge et al, 1997). In addition, proplatelet formation and polyploidization was observed (Fig.4C). Terminal differentiation of these cells in serum free media containing TPO/IL1b for 6 days (in total 17 days from iPSC colony differentiation initiation) led to an increase in cell size and appearance of polyploid cells (Fig.4D/E red arrows). Flow cytometry-based counting of cells that express CD41a, showed a megakaryoid yield of 6.9*10^3^ cells/iPSC (pool harvested between day 11-18) (Fig.4F). This number is significantly higher compared to traditional CD34^+^ megakaryoid expansion (1.5*10^1^/CD34^+^ cell). These results show that single cell-initiated differentiation directed to megakaryoid lineage successfully produce large quantities of relatively pure megakaryoid cells (~74%) that are capable to terminally differentiate towards mature megakaryocytes, and show the hallmarks of megakaryopoiesis (Bluteau et al, 2009; Geddis, 2010).

**Figure 4.**
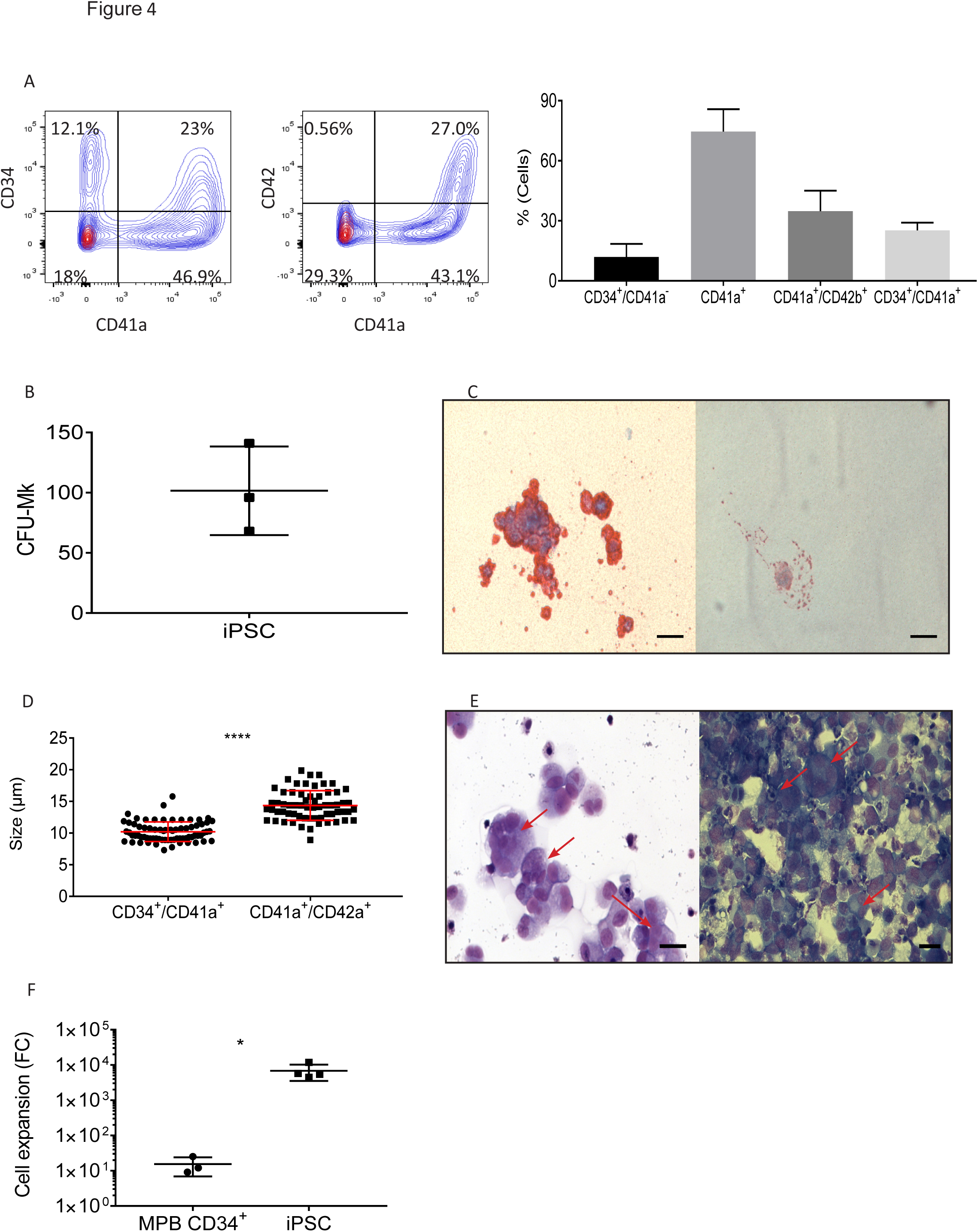
iPSC differentiation to megakaryocyte cells. **A)** Megakaryoid-associated marker expression of cells produced on day 11. Left: representative FACS plots (blue: staining, red: isotype). Right: FACS quantification graph represents average expression (n=4). **B)** CFU-Mk of iPSC-MKs from day 11 expanded for 6 days before seeding into megacult (n=3). **C)** CFU-Mk colony and proplatelet producing iPSC-MK (scale bar 20 μm). **D)** Quantification of size increase during maturation (day 11+6) by Imagestream on megakaryoblasts (CD34^+^/CD41a^+^) (73 cells) and megakaryocytes (CD41a^+^/CD42b^+^) (63 cells), p<0.0001 with ImageJ. **E)** Representative cytospin of HE stained terminal differentiating (day 11+6) iPSC-MKs, marked by red arrows (scale bar 20 μm). **F)** Quantification of MPB-CD34^+^ (n=3) and iPSCs (n=4) produced MK per input cell, p<0.05. “n” is biological replicates and all quantifications show mean and SD.

### iPSC differentiation to myeloid cells

Myeloid differentiation was initiated using a broad myeloid induction cytokine cocktail in serum-free media up to day 12 (figure 2A). The produced cells were further differentiated in serum-free media towards mature granulocytic cells with specific cytokines, among them the colony-stimulating factor (G-CSF), which constitutes the main driving force for the generation and terminal differentiation of granulocytes (Fig.2A) (Bronchud et al, 1988; Welte et al, 1985). After 7 days additional expansion (day 12+7), the cell population represented a 95% pure CD18^+^/EMR3^+^ myeloid population. Committed eosinophil- and neutrophil-like cells could be identified within this myeloid population, which was accompanied with appropriate segmented nucleus morphology (Fig.5A). This population was negative for CD34 and expressed surface markers from the myeloid lineage: CD15^+^ (55%), CD11b^+^ (77%), CD18^+^ (95%) and granulocytic lineage: EMR3 (96%; neutrophil marker/eosinophil), SIGLEC8 (76%; eosinophil marker), CD123 (60%; basophil marker; Fig.5B and Figure S3). On average, a single iPSC gave rise to ~2.5*10^3^ cells after 12 days of iPSC differentiation, which could be further expanded and differentiated for 7 days resulting in ~2*10^4^ cells/seeded iPSC (Fig.5C). The 7 days additional expansion/differentiation (day 12+7) gave rise to CD15^+^ suspension cells at a quantity of ~2.5*10^6^ cells/6 cm dish (Fig.5C). Importantly, this cell population did not express erythroid (CD36, CD235a), megakaryoid (CD42b), monocytic (CD14) or lymphoid (CD56, CD19) lineage markers, confirming granulocytic commitment and high-purity (Figure S3). Functionally, iPSC-derived granulocytes (day 12+7) produced reactive oxygen species (ROS) upon PMA stimulation, which is a hallmark of neutrophils and eosinophils, and the cells displayed peroxidase activity (Fig.5D) (Lacy et al, 2003). In addition, the suspension cells became more granular during the final stage of differentiation (day 12+7), affirming granulocytic commitment (Figure S4). Overall, the results show that single cell-derived iPSC differentiation directed to the myeloid linage is able to give rise to a broad but highly pure myeloid population that can be differentiated to a CD15^+^ cell population with multilobulated nuclei, which share functional similarities with mature neutrophilic granulocytes originating from blood.

**Figure 5.**
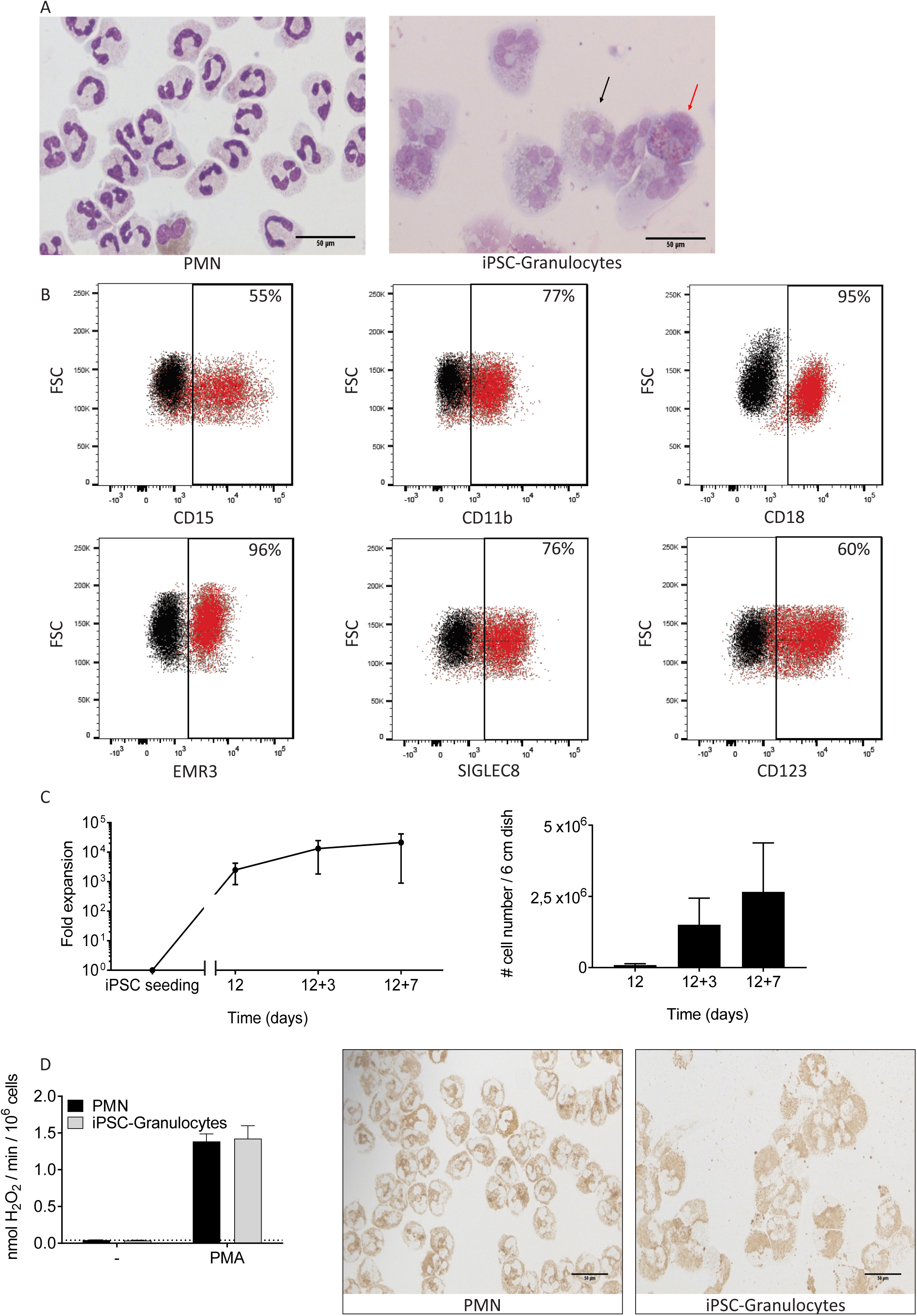
iPSC differentiation to myeloid cells. See also Figure S3, S4. **A)** Representative cytospin of neutrophils isolated from blood (PMN) (left) and iPSC-derived expanded cells at the final stage of differentiation (day 12+7) (right) after Giemsa-May staining (scale bar 50 μm). Black arrow; segmented nucleus. Red arrow; eosinophil-like cells. **B)** Myeloid/granulocyte-associated marker expression of expanded cells for 7 days (day 12+7) demonstrated by representative FACS plots (red: staining, black: isotype). **C)** Growth curve of expanded suspension cells/iPSC (Left). Amount of CD15^+^ cells during 7 days expansion (n=3). Quantities are given for 6 cm dish (Right). **D)** Left: production of reactive oxygen species (ROS) after activation with PMA in neutrophils isolated from blood (PMN) and in iPSC-derived granulocytes (day 12+7) (n=4). Right: representative cytospins of peroxidase-activity after benzidine staining of PMN or iPSC-derived granulocytes (day 12+7) (scale bar 50 μm). All quantifications show mean and SD, “n” is biological replicates.

## Discussion

iPSC differentiation provides a tool to study hematopoietic development, disease modelling/drug testing and holds a promise for future regenerative medicine e.g. the large-scale production of blood products. However, its potential application is hampered by the I) variations at (epi)genetic level between iPSC lines or between clones, II) incomplete understanding of differentiation cues, III) lack of standard (GMP-grade) protocols, IV) low cellular yield and V) low purity, often requires pre-purification. The iPSC culture and differentiation protocol we describe here, starts with a novel and easy to upscale single cell-seeding step, resulting in colonies with controlled number and uniform size, leading to a more reproducible system. Importantly, this system is feeder-free and avoids EB formation step further facilitating its application. In contrast to already available methods, here we describe an efficient way to produce three major blood cell types originated from a single differentiation protocol. When hematopoietic progenitors appeared, lineage-specific growth factor supplementation in serum free media matured these cells into well characterized erythroid, megakaryoid and myeloid cells. Our lineage specific conditions enabled us to produce specific blood cells up to ~95% purity, without any overexpression or pre-purification step, at large numbers which can be applied in hematopoietic developmental studies, and disease modelling/drug testing. However, currently the field is facing the difficulty to terminally differentiate towards end stage blood cell types, but most importantly to show the functionality of these cells, preventing clinical application at the moment.

In EB-based differentiation the hematopoietic cells are constrained within the 3D structure, hence it is difficult to isolate and analyse the efficiency of hematopoietic specification (Nii et al, 2015; Sivalingam et al, 2016; Sweeney et al, 2016). Co-cultures with feeder cells (e.g. OP9), are dependent on the feeder cell line batch, affecting reproducibility, and do not satisfy GMP requirements to facilitate future cellular therapy purposes. Salvagiotto et.al. (2011) were the first to introduce a single cell-seeded iPSCs differentiation in a feeder-free 2D system, but did not focus on terminal differentiation towards specific cell types or any form of large-scale production of the derived cells, which we show here (Salvagiotto et al, 2011). Niwa et.al. (2011) showed an alternative 2D approach using clump-passaged iPSCs, which is known to cause variation in colony size that may introduce a level of unwanted variability in culture outcome (Niwa et al, 2011). Here we demonstrated that a tight control of the starting conditions (e.g. single cell-seeding, colony number, colony size) could indeed lead to a robust, reliable differentiation towards hematopoietic progenitors and their terminal maturation to erythroid, megakaryoid and myeloid lineages, using a simple three step differentiation protocol. Importantly, the low single cell-passaging/seeding can be utilized for upscaling regardless of iPSCs lines.

Our iPSC differentiation protocol is also compatible with the use of vitronectin instead of matrigel, thus providing a completely xeno-free system (data not shown). In addition, this protocol is performed using GMP-grade Stemline II and our home-made culture media (named: Cell-Quin hereafter) which is GMP-grade, serum-free, defined and humanized (published by others, with minor modifications (Migliaccio et al, 2010)). Furthermore, avoiding co-culturing and the labour-intensive EB initiation step, we maintained a feeder-free system moving towards clinical application.

### iPSC differentiation to Erythrocytes

The potential of iPSCs differentiation to the erythroid lineage was immediately recognized and utilized in early blood developmental studies, disease modelling but also as a future source to produce transfusion ready red blood cells (Lapillonne et al, 2010). Several feeder- and serum-free methods have subsequently been developed, however most protocols are associated with low yield, low purity and suboptimal differentiation ability (Choi et al, 2009a; Dorn et al, 2015; Kobari et al, 2012; Smith et al, 2013). Here we show that monolayer differentiation starting from single cell-derived iPSC colonies produces CD71^+^/CD235^+^/CD36^+^ EBLs within 9-10 days. The combination of EPO, hSCF and glucocorticoids induces proliferation of immature pro-EBLs, inhibiting the induction of terminal differentiation and synchronizing erythroid cultures in the pro-EBL/basophilic stage (Bauer et al, 1999; Leberbauer et al, 2005; van den Akker et al, 2010; von Lindern et al, 1999). iPSC-derived cells could be expanded in this condition for at least 9 days (day 12+9) to pure 98.5% CD71^+^/CD235a^+^ EBLs and basophils without apparent terminal differentiation. These expanded iPSC-erythroid cells (day 12+9) could all be synchronously matured to late orthochromatic normoblasts within 7 days expressing 90.6% CD71^+^/CD235a^+^ and 7% CD71^-^/CD235a^+^ indicating reticulocyte formation (Socolovsky et al, 2001). Recently, Olivier et.al. reached similar results compared to ours (~0.5-2*10^5^ 99.1% (CD235a^+^) pure erythroid cell/iPSC), but within 31 days (including terminal differentiation). However their system is complex containing 7 steps including EB (3D) formation (Olivier et al, 2016). In comparison to other feeder-free monolayer protocols, Salvagiotto et.al. used a 31 days differentiation-scheme reaching mature erythroid cells expressing 38.6% CD71^+^/CD235a^+^ and 24.5% CD71^-^/CD235a^+^ with 30.7% CD71^-^/CD235a^-^ cells indicating a highly heterogeneous cell population (Salvagiotto et al, 2011). While Niwa et.al. defined erythroid population was 36% CD235a^+^ after 30 days of differentiation, but did not show the erythroid differentiation efficiency. Furthermore, no erythroid yield was shown (Niwa et al, 2011), which is generally underreported in published protocols. We show that differentiated iPSC-erythroid cells follow primarily a definitive wave of fetal erythropoiesis confirmed by the high expression level of fetal- and small portion of adult Hb. A consistently low amount of embryonic Hb was observed, corresponding to the presence of a small primitive erythropoiesis wave. These results meet with current literature demonstrating that pluripotent stem cell differentiations, irrespective of the used system, leads to embryonic/fetal definitive developmental stages (Chang et al, 2006; Dorn et al, 2015; Olivier et al, 2016; Yang et al, 2014). Within this definitive fetal wave we observed a small percentage of adult Hb in line with few other protocols (Lu et al, 2008; Sivalingam et al, 2016; Yang et al, 2014). The data suggests that ~75% of hematopoiesis through our 2D protocol is definitive, which may be extrapolated to megakaryoid and myeloid lineages. However, due to the absence of known fetal markers discriminating embryonic from definitive fetal/adult in the myeloid and megakaryoid lineages this remains difficult to assess.

### iPSC differentiation to Megakaryocytes

Megakaryoid differentiation from iPSC is hampered by low purity of megakaryoid cells and lack of expansion ability. Although, lentiviral-mediated over-expression of megakaryoid (transcriptional) regulators may be incompatible with cellular products due to genomic integrations or re-activation of exogenous genes, it has been successfully used to improve megakaryoid specification and to increase megakaryoblast expansion (Lim et al, 2013; Moreau et al, 2016; Nakamura et al, 2014; Takayama et al, 2010). Moreau et.al. used exogenous transcription factor expression of GATA1, FLI1 and TAL1 to generate pure MK with a yield of 1*10^5^ cells within a long culture period of 90 days using a EB-base protocol (Moreau et al, 2016). Nakamura et al used c-MYC, BMI1, and BCL-XL over-expression to increase survival and proliferation of megakaryoblasts, but was dependent on a C3H10T1/2 feeder layer (Lim et al, 2013). Interestingly, our 2D approach, without any further genetic modification or feeder layer, resulted in 7*10^3^ cells within 17 days and a mean megakaryoid purity of 74%. Qiang Feng et.al. was capable of producing ~16 MK/iPSC without any forced expression of transcription factors, which we could increase ~429 fold showing the importance of defining a robust iPSC differentiation initiating protocol (Feng et al, 2014). In addition, these megakaryoid cells have high CFU-Mk potential, with the ability to produce proplatelets. The combination of this system together with the introduction of megakaryoid regulators or small molecules in a footprint-free manner may further increase this yield.

### iPSC differentiation to myeloid cells

Most described myeloid differentiation protocols are 3D and/or apply co-culture with feeder cells, making these systems not ideal for future clinical applications (Brault et al, 2014; Sweeney et al, 2016; Zou et al, 2011, Lim, 2013 #1235). With the introduction of our 2D iPSC differentiation system, we were able to produce on average 2*10^4^ myeloid committed cells/iPSC, that was mostly consisting of multilobulated granulocytes and were 95% myeloid. Lachman et.al. reported a yield of 3-7*10^5^ granulocytes per 1 well/6 wells plate, whilst the 2D system described here yielded a higher production of myeloid cells/iPSC ratio (Lachmann et al, 2015). The studies of Morishima et.al. and Nayak et.al. focused on iPSC-derived neutrophil differentiation using a monolayer culture system (Morishima et al, 2014; Nayak et al, 2015). Both studies showed the generation of mature neutrophils with the presence of granule proteins such as ELASTASE, LACTOFERRIN or MYELOPEROXIDASE, and ROS production upon PMA stimulation, which we have also shown. Although, neither of these studies described the yield compared to input iPSCs, making it impossible to compare it with our system with respect to yield.

## Conclusions

In conclusion, the described 2D hematopoietic differentiation protocol represents a reliable system to generate committed erythroid, megakaryoid and myeloid cells of high-purity, without overexpression of transcription factors and/or pre-purification. Despite the significant yield of committed hematopoietic (progenitor) cells of different lineages reported here, the field of hematopoietic differentiation from iPSC is struggling to overcome the last steps of megakaryocyte and erythroid differentiation, namely platelet formation and enucleation/reticulocyte formation respectively. Indeed, also here we do not provide answers for these culture problems but introduce a robust and reproducible culture system to rapidly tackle these question using for instance overexpression of specific proteins or through performing drug screenings to identify compounds that are accommodating these last differentiation steps. In general, the described method can be used for large-scale production of cells for research and downstream applications.

## Acknowledgements

This study was supported by funding from Sanquin (MvL, EvdA, TK, CA and MH: PPOC: 15-2089), the Landsteiner Foundation (LSBR1141; EV and TW) and ZonMW (40-41400-98-1327). We would like to thank the Sanquin central Facility for flow cytometry/sorting expertise (Sanquin Central Facility, Amsterdam, the Netherlands).

